# Histological analysis of Scots pine (*Pinus sylvestris* L.) seedlings in response to root rot pathogen *Heterobasidion annosum* inoculation

**DOI:** 10.1101/2024.01.17.576084

**Authors:** Khaled Youssef, Salla Marttila

## Abstract

*Heterobasidion* root rot is one of the most serious and economically destructive forest diseases in the Northern Hemisphere. Although several studies have explored the genetic and chemical responses of Scots pine to *Heterobasidion* spp. infection, the histological defense responses of this species remain poorly understood. In this study, we investigated the histological responses of three-year-old Scots pine seedlings to *Heterobasidion annosum* inoculation and a wounding treatment with no inoculation, focusing on lesion length and traumatic resin-duct characteristics (density, size). Our results showed that *H. annosum*-inoculated seedlings exhibited significantly more browning necrotic lesions than wounded seedlings. Traumatic resin duct density was significantly higher in *H. annosum*-inoculated seedlings compared to wounded seedlings, particularly within the first two cm from the inoculation point. However, as the distance from the inoculation point increased, the resin duct density decreased. Notably, there were neither statistically significant differences in the mean size of traumatic and constitutive resin ducts between the two treatments, nor within the *H. annosum* inoculation treatment itself. In contrast, within the wound treatment, the mean size of traumatic resin ducts was found to be significantly smaller than that of constitutive resin ducts. Furthermore, traumatic resin ducts did not prove to be reliable markers for dating *H. annosum* infection in Scots pine. Overall, this study advances knowledge about Scots pine’s histological defense mechanisms against *H. annosum* invasion, which has significant implications.

## 1 Introduction

Scots pine (*Pinus sylvestris* L.) like other conifers, has developed multiple constitutive and inducible defense mechanisms to protect itself from a wide range of biotic and abiotic stressors (Krokene 2015; Franceschi et al. 2005). Constitutive defenses including such as parenchyma cells filled with phenolics (Franceschi et al., 2000; Krekling et al., 2000), and resin ducts containing oleoresin (hereinafter, resin) (Wu and Hu, 1997), form the first defense line and are produced in the absence of any attack or disturbances. However, when the tree is attacked by insects or pathogens, the inducible defenses such as increasing resin and formation of traumatic resin ducts, are activated to kill the invaders or restrict their further expansion within the tree (Franceschi et al. 2005; Vázquez-González et al. 2020). A significant defense mechanism of conifer trees to fungal and insect attacks is the synthesis of resin which is a complex mixture of various volatile monoterpenes, sesquiterpenes, and diterpenes, and possess antifungal, antibacterial, and anti-insect properties (Celedon, 2019; Keeling, 2006). Resin is synthesized, stored, and transported within a complex network of resin ducts (resin canals). Resin ducts are multicellular tube-like structures composed of secretory epithelial cells that synthesize and release resin into the ducts’ lumen, where it is preserved under pressure (figure 1, A) (Krokene and Nagy, 2012; Nagy et al., 2000). Based on their arrangement, resin ducts are classified into two distinct types: axial ducts and radial ducts. Axial ducts are aligned parallel to the tracheids and more common in the xylem, while radial ducts, run through the rays in a radial direction and more abundant in the phloem (Beck, 2010; Wu and Hu, 1997). A large network of connections Scots pine (*Pinus sylvestris* L.), like other conifers, has developed multiple constitutive and inducible defense mechanisms to protect against a wide range of biotic and abiotic stressors (Krokene 2015; Franceschi et al. 2005). Constitutive defenses such as parenchyma cells filled with phenolics (Franceschi et al., 2000; Krekling et al., 2000), and resin ducts containing oleoresin (hereinafter resin) (Wu and Hu, 1997), constitute the primary defensive barrier and are produced in the absence of any attack or disturbance. However, when a tree is attacked by insects or pathogens, inducible defenses such as increasing resin and forming traumatic resin ducts are activated to kill the invaders or restrict their further expansion within the tree (Franceschi et al. 2005; Vázquez-González et al. 2020). For conifers, a significant defense mechanism against fungal and insect attacks is the synthesis of resin which is a complex mixture of various volatile monoterpenes, sesquiterpenes, and diterpenes, and which has anti-fungal, anti-bacterial, and anti-insect properties (Celedon, 2019; Keeling, 2006). Resin is synthesized, stored, and transported within a complex network of resin ducts (resin canals). Resin ducts are multicellular tube-like structures composed of secretory epithelial cells that synthesize and release resin into the ducts’ lumen, where it is preserved under pressure (figure 1, A) (Krokene and Nagy, 2012; Nagy et al., 2000). Resin ducts are classified based on their arrangement as either axial duct or radial ducts. Axial ducts are aligned parallel to the tracheids and more common in the xylem, while radial ducts run through the rays radially and are more abundant in the phloem (Beck, 2010; Wu and Hu, 1997). A large network of connections between the radial and axial resin ducts forms a complex resin reservoir, allowing resin to flow both vertically along the trunk and horizontally towards the bark’s surface (Figure 1, B) (Franceschi et al., 2005; Nagy et al., 2006).

**Figure 1.**
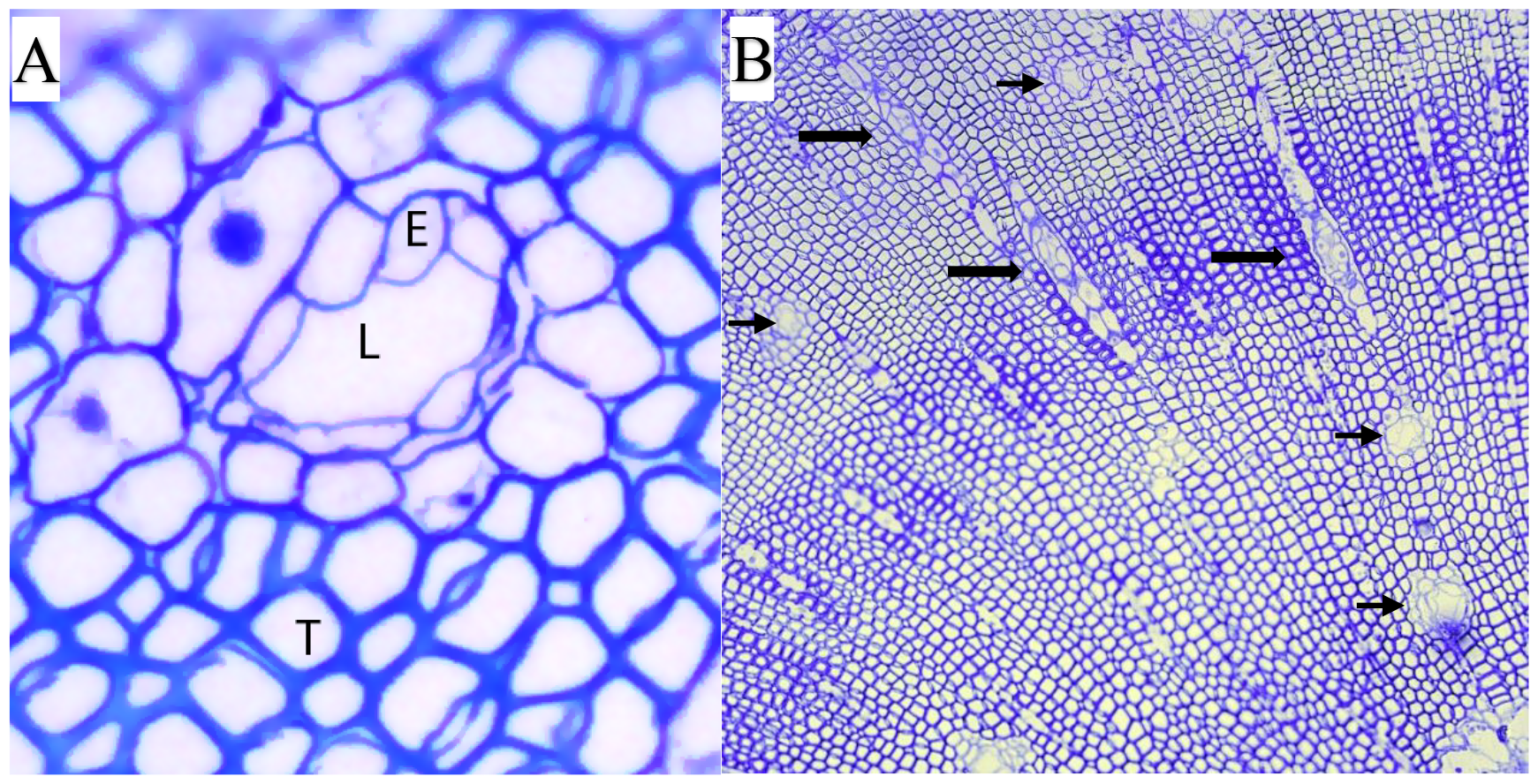
Transverse sections of the stem of a three-year-old Scots pine seedling. A) The structure of resin ducts; E: Epithelial cells, L: lumen, T: tracheids. B) Axial resin ducts (small arrows) and radial resin ducts (large arrows).

Functionally, resin ducts have two types: constitutive resin ducts (CRDs) and induced or traumatic resin ducts (TRDs). CRDs are dispersed throughout the woody tissues of the tree and their abundance varies among conifer species. They are relatively less abundant in the genera *Picea, Abies* and *Pseudotsuga*. In contrast, pine trees (*Pinus* spp.) and particularly Scots pine have a highly-developed network of CRDs (Wu and Hu, 1997). On other hand, traumatic resin ducts (TRDs) are typically produced in densely packed, compact bands, displaying distinct distribution patterns (Franceschi et al., 2005; Wu and Hu, 1997). They form as a defensive response to a wide array of both abiotic and biotic stress factors, including mechanical wounds (Gärtner and Heinrich, 2009; Luchi et al., 2005; Nagy et al., 2000), insect infestations (Deflorio et al., 2009; Franceschi et al., 2005; Krekling et al., 2000; Nagy et al., 2000), pathogenic fungal invasions (Krokene et al., 2003; Luchi et al., 2005; Mercado et al., 2023), as well as exogenous application of methyl jasmonate (Heijari et al., 2005; Krokene et al., 2023; López-Villamor et al., 2021; Martin et al., 2002).

Investigations on Norway spruce and Douglas-fir have demonstrated that both physical injury and fungal infections can trigger the development of traumatic resin ducts in tissues one meter from the initial induction site (Cruickshank et al., 2006; Krekling et al., 2004). Several earlier studies have demonstrated a connection between the quantity and structures of resin ducts and their role in enhancing tree resistance to various insects and pathogens likely through increasing resin synthesis (Krokene et al., 2003; Schmidt et al., 2011; Zhao and Erbilgin, 2019) and inhibiting the inward expansion of the invaders (Krokene and Nagy, 2012; Nagy et al., 2000).

Alongside their role in enhancing conifers’ resistance to several biotic stressors, traumatic resin ducts have been used as a tool to date previous tree damage events, such as geomorphic events (Schneuwly et al., 2009; Stoffel, 2008), bark beetle outbreaks (Derose et al., 2018), and *Armillaria* root rot infection in Douglas-fir (Cruickshank et al., 2006).

Root rot caused by *Heterobasidion* spp. is one of the most serious and destructive diseases of conifer forests in the northern hemisphere (Asiegbu et al., 2005; Garbelotto and Gonthier 2013). The pathogen enters a stand by airborne basidiospores that infect freshly-cut stump surfaces and then spread vegetatively to the neighboring living trees through root contacts (Rishbeth 1951; Asiegbu et al., 2005; Garbelotto and Gonthier, 2013).

Infected mature pine trees appear externally healthy for decades (Kurkela, 2002; Wang et al., 2014). However, externally-obvious signs, such as crown defoliation and the exudation of resin, only become apparent in the late stages of infection (Asiegbu et al., 2005b; Kurkela, 2002). These external symptoms are not reliable markers of *Heterobasidion* spp. infection nor do they determine the time of infection (Kurkela, 2002; Rönnberg et al., 2006). Several recent studies have investigated the genetic and chemical responses of Scots pine to *Heterobasidion* spp. infection and highlighted the role of candidate genes within the terpenoid pathway in Scots pine’s resistance to *H. annosum* (Mukrimin et al., 2019). Additionally, they observed that Scots pine genotypes resistant to *Heterobasidion* infection exhibited higher levels of 3-carene, while susceptible genotypes exhibit elevated concentrations of α-pinene (Liu et al., 2022b). Nevertheless, the histological defense responses of Scots pine to *Heterobasidion* spp. have received limited exploration. It is unclear whether the inducibility and formation of TRDs observed in Norway spruce will likewise occur in pine.

Better understanding of the defense mechanisms employed by Scots pine against *Heterobasidion* infection is crucial for improving the production of superior plant materials, thereby mitigating the impact of this disease.

The primary objective of this study is to provide a quantitative analysis of the inducible histological defense responses observed in the xylem of Scots pine seedlings in response to *Heterobasidion annosum* inoculation. Specifically, this study aims to:

- Quantify the length of necrotic lesions that develop in response to *Heterobasidion annosum* inoculation and wound treatment.
- Characterize traumatic resin ducts in terms of their distribution, density, and size as a result of both *H. annosum* inoculation and wound treatment.
- Investigate the feasibility of using traumatic resin ducts as a marker to determine the timing of infection.

## 2 Materials and Methods

### 2.1 Plant materials

In this study, apparently-healthy and vital three-year-old bare-rooted Scots pine (*Pinus sylvestris*, L.) seedlings from a commercial nursery (Ramlösa Plantskola AB, Sweden) were used. The geographical origin of the seed material was Målilla, Småland, Sweden (seed batch family code: S94/1082). On June 1, 2018, the seedlings were transplanted into 5 L pots and afterwards placed in a greenhouse located at the Alnarp campus of the Swedish University of Agricultural Sciences. All seedlings were subjected to ambient lighting and temperature conditions and watered twice per week for the duration of the experiment.

### 2.2 Fungal material and inoculation

*Heterobasidion annosum* (isolate B11) isolated from Scots pine trees by LiYing Wang and Jonas Rönnberg in 2012 was used in this study. Prior to inoculation, the fungal strain was identified by species-specific primers (Hantula and Vainio, 2003).

The *H. annosum* isolate was cultured on Hagam agar and grown at room temperature for three weeks before the inoculation, which was done on June 21, 2018. Inoculation points were approximately 6 cm above the soil level. Each stem was wiped with 70% ethanol, then the bark was carefully removed in a rectangular configuration like a window, using a sterile scalpel to access the xylem surface. Subsequently, 5 mm diameter round plugs, derived from the actively-developing *H. annosum* culture, were placed onto the exposed surface and sealed with Parafilm. For the wounding treatment (mock-inoculation), sterile Hagam agar plugs without fungus were used.

### 2.3 Data sampling

Six months after inoculation, all seedlings were harvested. The stem of each seedling was cut in half symmetrically along the stem axis (longitudinally) to observe necrosis in the xylem. After that, the stems were cut at five 1 cm intervals upwards of the inoculation.

#### 2.3.1 Sample preparation for light microscopy

Samples were prepared for light microscopy as in Ghasemkhani et al., 2016. Briefly, the samples were fixed by immersing them in a mixture of 2.5% (v/v) glutaraldehyde and 2% (w/v) paraformaldehyde in 0.1 M Na-phosphate buffer, pH 7.2, with gentle shaking for 4 h at room temperature, followed by 24 h at 4 °C. Then, the samples were washed with 0.1 M Na-phosphate buffer for 3 × 15 min and stored at 4 °C in the buffer until the next step. After washing the fixed samples with distilled water for 3 × 15 minutes, they were dehydrated with acidified 2-dimethoxypropane (DMP) for 2 × 30 minutes and acetone for 2 × 20 minutes. For infiltration with plastic resin, fresh Spurr’s resin (low viscosity kit; Ted Pella, Redding, CA, USA) was prepared according to the manufacturer’s instructions for infiltration with acetone-resin (3:1) for 3 h, (1:1) for 3 h, and (1:3) for 24 h, then 100% resin for 24 h, followed by 6 h in fresh 100% resin the next day. All steps were carried out in a fume hood with gentle shaking. Polymerization was performed with fresh resin in polyethylene capsules at 70 °C for 24 h, followed by 48 h at 40 °C, and post-polymerization in a fume hood a few days before sectioning. Cross sections (1-5μm) were cut with an ultramicrotome and placed on adhesion slides (Superfrost ® Plus, Menzel-Gläser, Braunschweig, Germany) for light microscopy.

### 2.4 Light microscopy

To observe general tissue structure, Toluidine Blue O (Sigma-Aldrich, Steinheim, Germany) was used. The sections were stained for 30 seconds on a hot plate with filtered 1% Toluidine Blue O in 1% Na-tetraborate, rinsed with water, air-dried, and mounted with Pertex ®. Sections were studied with a Leica DMLB light microscope, and images were taken with a Leica DFC450C digital camera (Leica Microsystems) at low magnification (5x objective).

### 2.5 Necrotic lesion length measurement

The tissue-level induced defense responses were determined by necrotic lesion length and resin ducts characteristics. The necrotic lesion length in the xylem was measured along the plant stem using a digital ruler.

### 2.6 Resin duct measurement

The resin duct system within stems of pine seedlings was assessed by examining the following characteristics: the density of traumatic resin ducts per unit area of the measured area of the last annual growth ring (TRD/mm^2^), and the size of traumatic resin ducts (μm^2^) in the last annual growth ring, as compared to the constitutive resin ducts (CRDs) in the previous year’s growth ring. The measurements were performed at distances of 1-5 cm in 1 cm intervals from the inoculation point or wound. The size (area) of resin ducts, including the lumen and the epithelial cells, was measured using the ImageJ image-processing program (Abràmoff et al., 2004).

### 2.7 Statistical analysis

T-tests for independent samples were used to assess the statistical differences in necrotic lesion length between the two treatments, and for differences between the mean size of traumatic resin ducts and constitutive resin ducts within treatments. Linear mixed models using the lmer() function from the *lme4* package in R were used to analyze the impact of treatments and distance from inoculation point on the density and mean size of traumatic resin ducts. Treatments and distance from the inoculation point were included as fixed factors while tree and resin duct types were included as random factors (Vázquez-González et al., 2019).

## 3 Results

Statistically significant treatment effects were found out for most variables, although the number of replicates was limited.

### 3.1 Necrotic lesion length

Scots pine seedlings inoculated with *Heterobasidion annosum* produced significantly greater browning necrotic lesions compared to the wound treatment (unpaired two-sample t-test; p=0.000002, n=9; Figure2).

**Figure 2.**
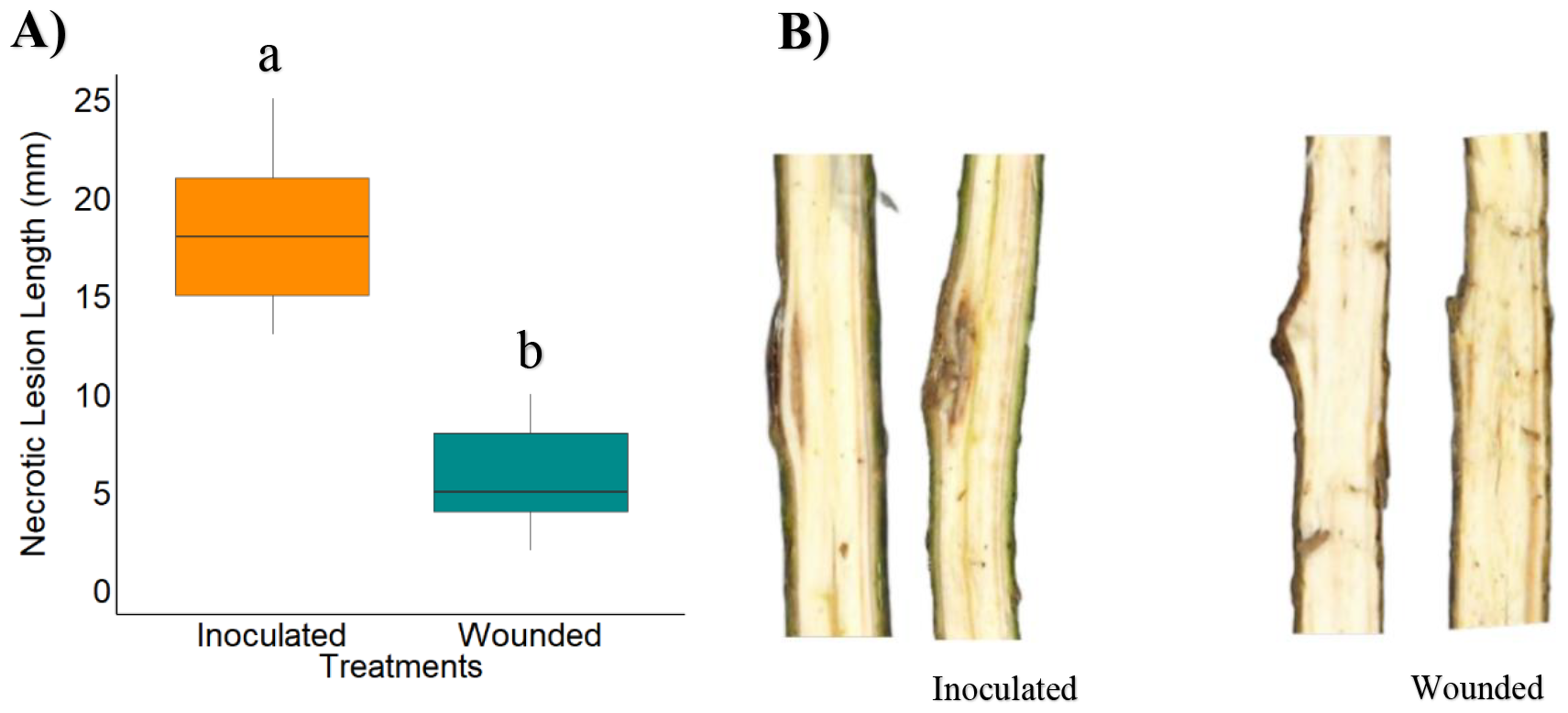
A) The length of necrotic lesions (mm) in the xylem of Scots pine seedlings in response to *H. annosum* inoculation and wound treatment. Different letters indicate significant differences at p<0.05 between the two treatments. B) Necrotic lesions seen in the xylem of two representative Scots pine seedlings in response to both treatments.

#### Characteristics of traumatic resin ducts

Resin ducts could easily be seen in the woody tissue under low magnification (Figure 3). Traumatic resin ducts seemed to be distributed in a more organized way as compared to the constitutive resin ducts which were more unevenly distributed. The mixed effect model showed that the density of traumatic resin ducts in *H. annosum*-inoculated seedlings was significantly higher compared to the wound treatment (p=0.036, n=3; Table 1, Figure 3). The differences were most pronounced closer to the inoculation point (≤3 cm). However, as the distance from the inoculation point increased, the density of traumatic resin ducts decreased, and the differences were no longer statistically significant at distances of ≥4 cm (Figure 4).

**Figure 3.**
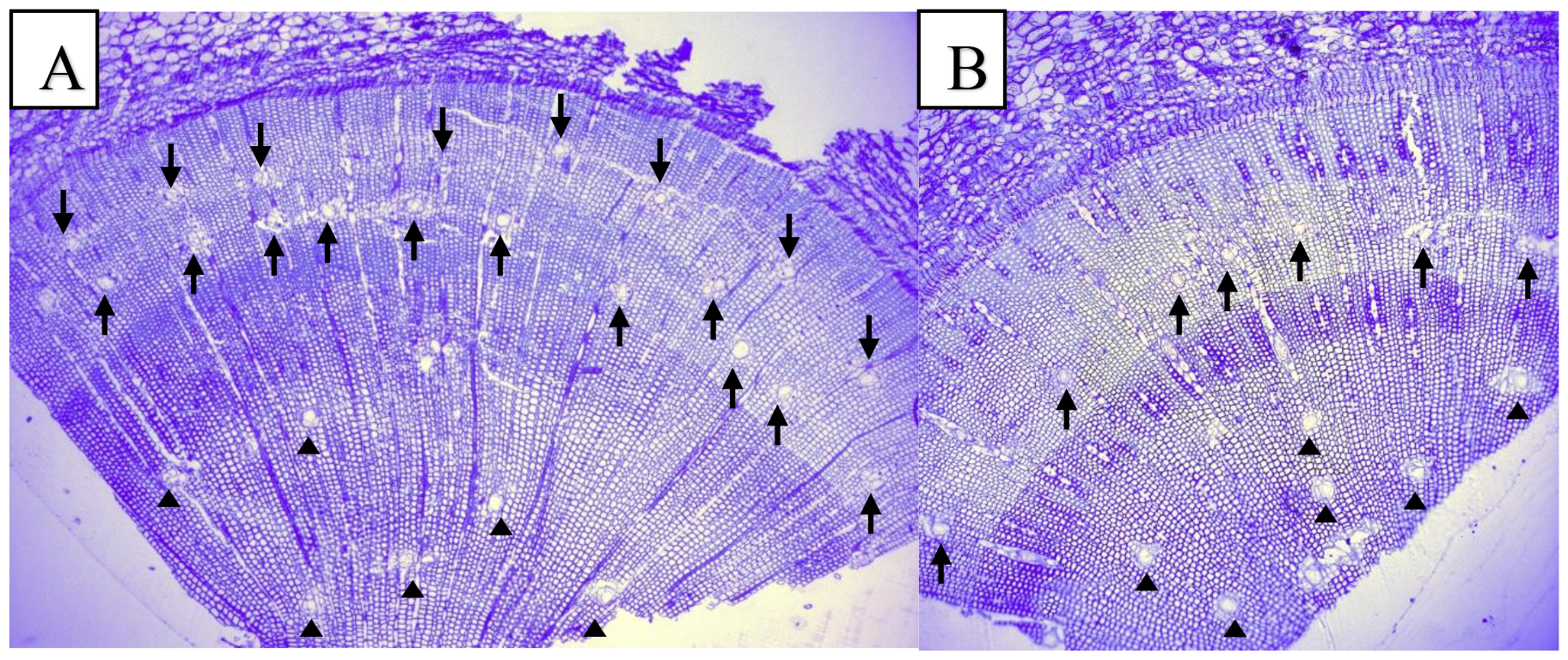
Resin-based defense response of three-year-old Scots pine seedlings to *H. annosum* (A) and wounding (B) treatments. In the last annual growth ring one or two rows of traumatic resin ducts (small arrowheads) are seen, while in the previous year’s growth ring only constitutive resin ducts are present (triangles).

**Figure 4.**
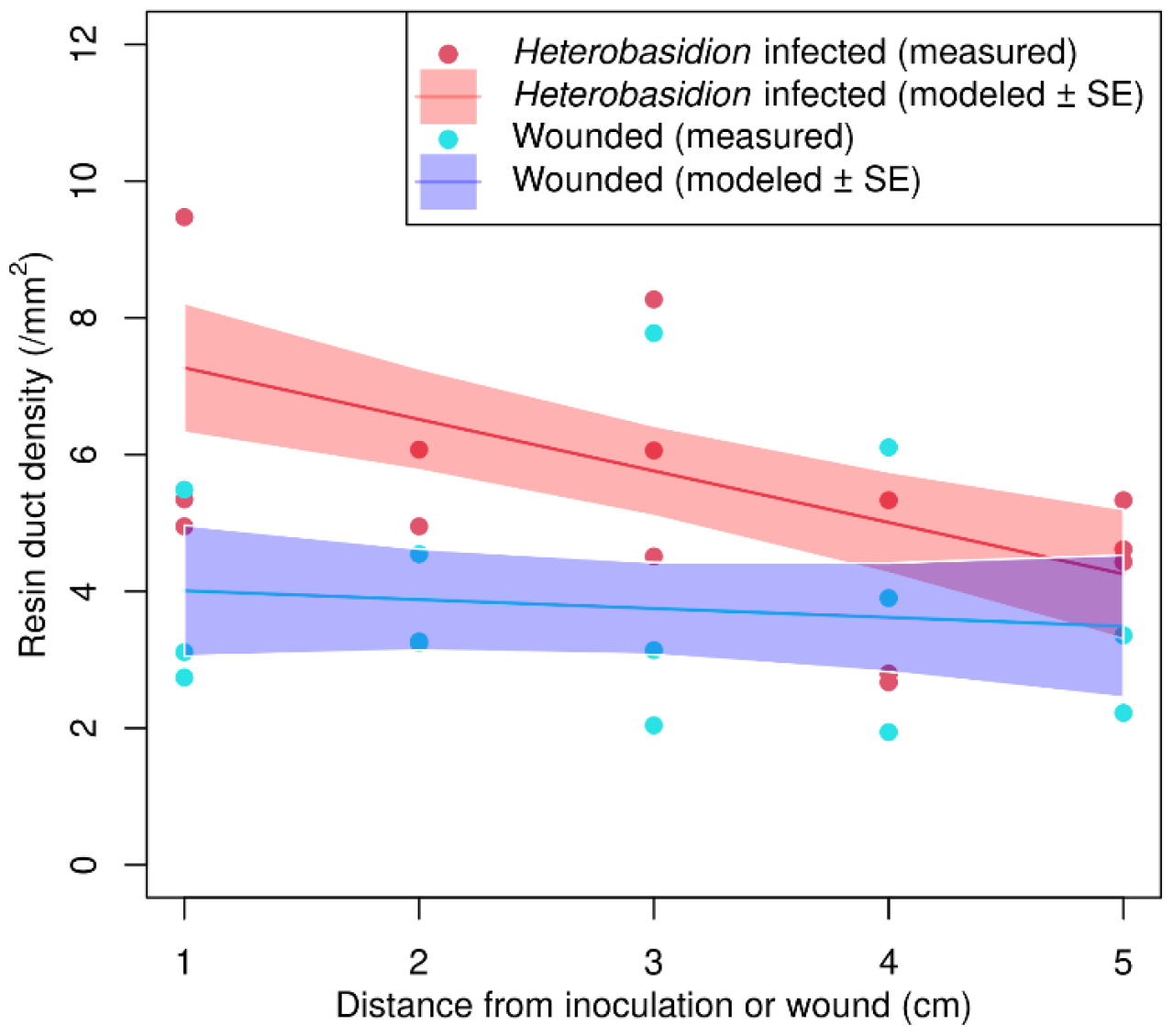
The resin duct density in Scots pine seedlings was evaluated following inoculation with *Heterobasidion annosum* and wound treatment at varying distances from the inoculation point. Individual seedling measurements are represented by data points, while the trend lines illustrate the mean responses estimated through the model. The shaded regions delineate the standard error associated with these mean estimates. Notably, the absence of overlap in standard errors up to 3 cm signifies statistically significant differences at a significance level of p<.05 within these specific distances from the inoculation point.

The mean size of traumatic and constitutive resin ducts did not differ significantly between *H. annosum*-inoculated and wound treatments (p-value: 0.51, p-value: 0.91 respectively). Furthermore, there were no significant differences between the mean size of traumatic resin ducts and constitutive resin ducts within the *H. annosum* inoculation treatment (p-value: 0.62). In contrast, within the wound treatment, the mean size of traumatic resin ducts was significantly smaller than that of constitutive resin ducts (p-value: 0.094) (Figure 5).

**Figure 5.**
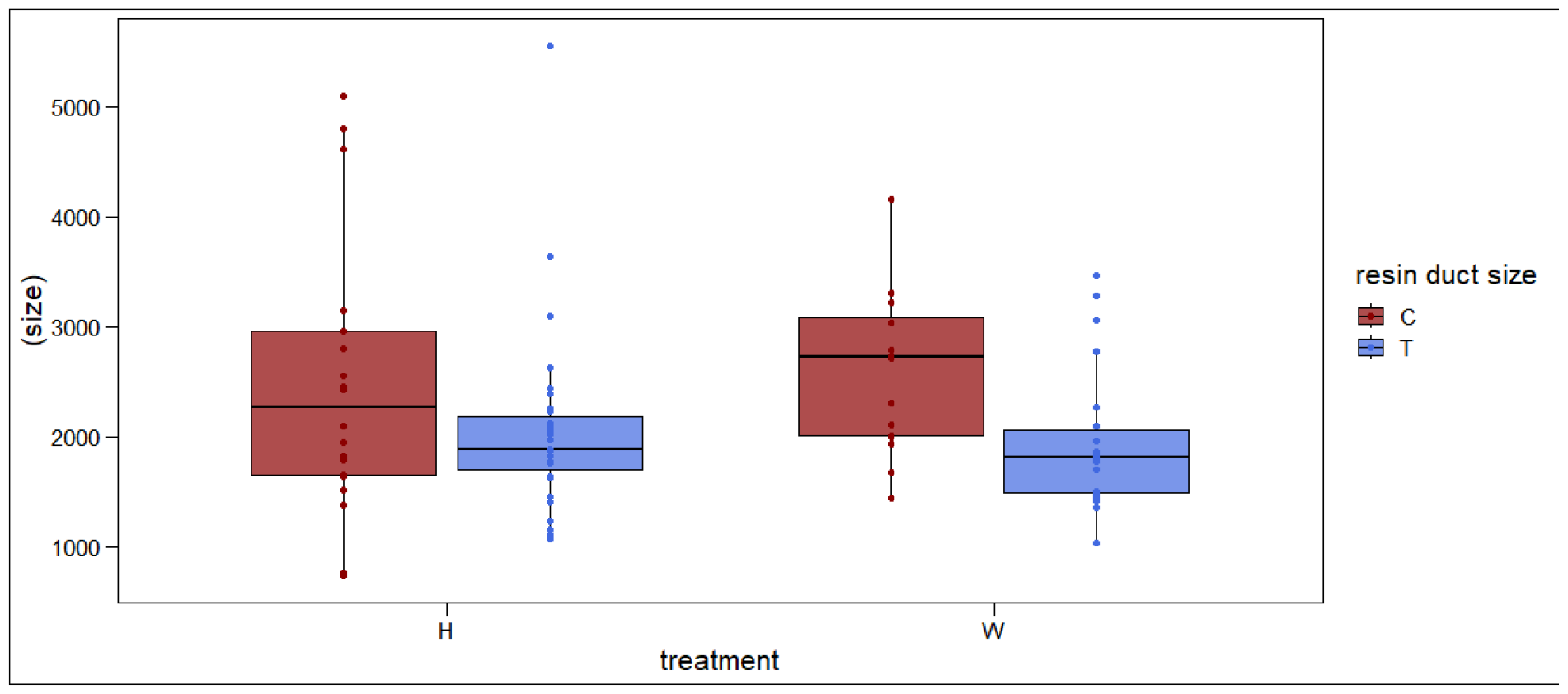
The mean size of traumatic resin ducts (μm^2^) (T) and constitutive resin ducts (C) for the two treatments: *H. annosum* inoculation (H), and wound treatment (W).

## 4 Discussion

In this study, we investigated and contrasted the anatomical responses of Scots pine seedlings to inoculation with the Heterobasidion root rot pathogen and wounding treatment. Our results showed that *H. annosum*-inoculated seedlings exhibited significantly greater browning necrotic lesions than wounded seedlings which is in accordance with precent studies (Johansson et al 2004; Mukrimin et al. 2019; Terhonen et al. 2019; Liu et al. 2020 a; Liu, Wang, Ghimire, et al. 2022). This difference can be attributed to the initial induced responses of pine seedlings to the pathogen invasion. This response aims to hinder the spread of the pathogen by decreasing the levels of sugars and while simultaneously accumulating potentially toxic compounds such as terpenoids, lignin, and phenolics in the vicinity of the inoculation site (Johansson et al., 2004; Liu et al., 2022a; Mukrimin et al., 2019; Raffa and Smalley, 1995). Besides, the ability of *H. annosum* to induce swiftly expanding necrotic lesion inside the xylem tissue of their hosts (Pearce, 1996). the increased size of lesion expanding, and the development of browning discoloration lesions serve as robust indicators of the pathogen’s successful colonization (Krokene and Solheim, 1998; Liu et al., 2020; Pearce, 1996). Necrotic lesion size (length) has frequently been utilized as marker for host resistance or susceptibility(Danielsson et al., 2011; Liu et al., 2022a; Mukrimin et al., 2019, 2018). The length of necrotic lesions observed in this study ranged from 13-25 mm, suggesting that each individual Scots pine seedling exhibited a variable level of susceptibility to *H. annosum*, i.e., seedlings with longer lesion lengths are considered as being more susceptible compared to those with shorter lengths. Therefore, the measurement of necrotic lesion length might serve as a marker during the initial phase of assessing and selecting Scots pine genotypes that may have significance in resistance breeding programs.

Resin ducts serve as a first anatomical and chemical defense mechanism, providing protection to conifer plants against insects and diseases. Following an attack or mechanical wound, the constitutive resin is transferred from resin ducts to the affected region in order to effectively seal the wounds and hinder the entry of harmful organisms (Berryman, 1972). The results of this study revealed that pine seedlings inoculated with *H. annosum* exhibited a greater density of traumatic resin ducts in comparison to the seedlings subjected to wounding treatment. It is hypothesized that this response serves as a mechanism utilized by seedlings to minimize the presence of unprotected gaps in the attack site, thereby restricting the further advance of the fungal hyphae and eventually killing it (Nagy et al. 2000). Furthermore, the higher the density of resin ducts, the higher the flow of resin (Ayres and Lombardero, 2000; Blanche et al., 1992). Similar pattern has been observed in Norway spruce (*Picea abies*), where spruce trees exhibited a higher density of traumatic resin ducts in response to *H. annosum* inoculation compared to sterile inoculation (Krekling et al., 2004). Lim et al. (2021) examined the stress response of five-year-old Scots pine xylem to mechanical wounding using RNA sequencing. and found that pine resin biosynthesis was not induced in response to wounding (Lim et al., 2021). Other authors have observed a higher concertation of terpenes in resistant pine trees against *H. annosum* than in susceptible ones (Liu et al., 2022b; Mukrimin et al., 2019). Furthermore, Gibbs, 1968 found that the resistance of pine trees to Heterobasidion root rot is correlated with their capacity to mobilize resin. Accordingly, our results showed that both resin and resin ducts play a significant role in enhancing pine resistance to Heterobasidion disease.

Conifer trees resistant to insects and pathogens are hypothesized to allocate a higher proportion of their resources to defense mechanisms. This hypothesis is supported by several studies which have demonstrated that resin duct characteristics, such as size and density, can be employed as indicators for biotic resistance and serve as valuable metrics of defensive investment (Christiansen et al., 1999; Ferrenberg et al., 2014; Kane and Kolb, 2010; Zhao and Erbilgin, 2019). For example, among pine species, those that are resistant to the mountain pine beetle invested more resources in resin duct defense than in growth, compared to susceptible species (Bentz et al., 2017). In this study, we found that wounded pine seedlings produced significantly less dens resin ducts compared to *H. annosum* inoculation and smaller in size compared to the constitutive ones, suggesting that Scots pine seedlings possess the capability to modulate the intensity of its defense responses in accordance to the severity of the presented challenges (Nagy et al, 2006).

In addition to their role in enhancing resistance to biotic disturbances, traumatic resin ducts have been employed as a marker for dating past events, since they persist in the wood throughout the tree’s lifespan, rendering them an annually resolved archive of defensive allocation for retrospective analysis (Hood et al., 2020). Cruickshank et al. (2006) employed traumatic resin ducts as a discernible indicator for dating *Armillaria ostoyae* within naturally infected Douglas-fir trees, detecting them from distances exceeding one meter away from identified lesions.

Derose et al., 2018 used the presence of traumatic resin ducts (TRDs) in tree rings to track spruce bark beetle outbreaks. They found that the number of survival trees with TRDs were much higher during the outbreak than it had been at any other point in the trees’ lives. Although our data revealed a significantly higher density of TRDs in *H. annosum* inoculation compared to wounding treatment, this significant response was localized and disappeared beyond three cm from inoculation point (figure 4). Furthermore, the strength of this response gradually diminished as the distance from the inoculation point increased for both treatments. Consequently, TRDs were not found to be a reliable marker for dating *H. annosum* infection in Scots pine. The local response of Scots pine to the both treatments is probably due to rapid containment of the fungal colonization and greater reliance on the constitutive anatomical mechanisms (Cruickshank et al 2006; Nagy et al 2006).

## 5 Conclusions

Our study provides additional insights into the histological defense mechanisms activated by Scots pine during *H. annosum* infection. We observed that Scots pine seedlings exhibit dynamic defense mechanisms with significant differences in necrotic lesion size and the characteristics of traumatic resin duct. These differences underscore the ability of Scots pine to modify its defense mechanisms depending on the specific type of threat it encounters. Traumatic resin ducts, however, did not prove to be a reliable indicator for dating the *Heterobasidion* infection in Scots pine. The demand for identifying alternative reliable markers for dating *Heterobasidion* infection in Scots pine and other conifers therefore remains a critical research focus.

## 7. Acknowledgments

We would like to thank Jonas Rönnberg and Michelle Cleary for their valuable contributions in the initial stage of this study. This research was funded by FORMAS, grant number 2015-01413; and the Rattsjö Foundation.

## Parsed Citations

